# Rbm24 maintains survival of cochlear outer hair cells by repressing Insm1

**DOI:** 10.1101/2024.06.26.600754

**Authors:** Chao Li, Luyue Wang, Shuting Li, Jie Li, Ying Lu, Zhiyong Liu

## Abstract

The inactivation of Rbm24, an RNA-binding protein, results in the degeneration of cochlear outer hair cells (OHCs) during the postnatal period. However, the specific molecular mechanisms underlying this OHC death remain elusive. To address this, we conducted a comprehensive analysis comparing the gene profiles of wild-type OHCs to those lacking Rbm24 (*Rbm24^-/-^)* at postnatal day 7 (P7). Our results revealed that the overall differentiation program of OHCs is delayed in the absence of Rbm24. Furthermore, the expression of Insm1, a crucial factor for OHC development that is normally switched off by P2, remains prolonged in *Rbm24^-/-^* OHCs. Interestingly, when Insm1 is overexpressed, it also leads to OHC death. Significantly, the OHC degeneration is much less severe when both *Rbm24* and *Insm1* are simultaneously inactivated. These findings shed light on the important role of Rbm24 in repressing Insm1 and its impact on OHC differentiation and survival. Our study provides valuable insights into the complex genetic signaling pathways involved in OHC development.

## INTRODUCTION

Two subtypes of sound receptor hair cells (HCs) are housed in the mouse auditory epithelium, also known as the organ of Corti: inner HCs (IHCs) and outer HCs (OHCs) (1). Adjacent to IHCs and OHCs are different subtypes of supporting cells (SCs) (2). Atoh1 is a master transcription factor (TF) in development of both IHCs and OHCs, and no HCs form in the *Atoh1^-/-^*mice (3). Severing as sound amplifiers, OHCs uniquely express the motor protein Prestin (encoded by *Slc26a5*) (4, 5). In contrast, IHCs act as the primary receptors and form ribbon synapses with the spiral (auditory) ganglion neurons (6–8). IHCs specifically express Fgf8, vGlut3 (encoded by *Slc17a8*), and Otoferlin (9–13). Three key TFs are known to be involved in determining whether cochlear sensory progenitors adopt IHC or OHC fate: Tbx2, Insm1 and Ikzf2 (1). Tbx2 is required in IHC fate specification, differentiation, and fate maintenance at adult ages, and *Tbx2^-/-^* IHCs would transform into OHC-like cells (14–16). Oppositely, OHCs overexpressing Tbx2 would transdifferentiate into IHC-like cells (15–17). Moreover, Atoh1 and Tbx2 together can reprogram neonatal SCs into IHC-like cells (14, 18). Different from Tbx2, Insm1 and Ikzf2 are expressed in OHCs, but not IHCs (19–22), and the OHC fate cannot be maintained in Insm1 or Ikzf2-deficient OHCs (19, 20, 22), and half of the *Insm1^-/-^* OHCs tend to become IHC-like cells (19, 20). Note that Insm1 is transiently expressed in nascent differentiating OHCs and become undetectable in OHCs by postnatal day 2 (P2) (20, 21).

Both HC subtypes, especially the OHCs, are vulnerable to genetic mutations or other ototoxic factors, and degeneration of IHCs or OHCs result in severe hearing impairment. RNA binding motif Protein 24 (Rbm24) is an RNA binding protein and is maintained in both IHCs and OHCs once it is turned on at ∼E15 (23). Currently, the mechanisms underlying degeneration of *Rbm24^-/-^*OHCs remains completely elusive. Starting with the single cell transcriptomic assay between wild type (WT) and *Rbm24^-/-^* HCs, the differentially expressed genes (DEGs) were uncovered. Our data suggested that the global differentiation program of OHCs was delayed, which was further evidenced by the defective hair bundle development. Intriguingly, expression of Insm1, which is the nascent OHC marker, is dramatically prolonged and maintained in the *Rbm24^-/-^*OHCs by P14. It suggests that Rbm24 is needed to repress Insm1 in postnatal OHCs. Moreover, overexpressing of Insm1 also resulted in a similar OHC degeneration phenotype. Finally, we demonstrated that additional deletion of Insm1 mitigated the cell death of *Rbm24^-/-^*OHCs. Collectively, our study provided new insights of how Insm1 is repressed in postnatal OHCs and revealed that reactivation of Insm1 expression is one of the key mechanisms underlying cell death of the *Rbm24^-/-^* OHCs.

## RESULTS AND DISCUSSION

### Single cell transcriptomic analysis of wild type and *Rbm24^-/-^* OHCs and IHCs

We aimed to decipher the global gene perturbations in the *Rbm24^-/-^*HCs, relative to the control (Ctrl) HCs at P7. According to our previous study, *Rbm24^-/-^* HCs have not started to degenerate at P7 (24). The Ctrl_HCs were manual picked via *Atoh1^P2A-Tdtomato/+^* mice in which both IHCs and OHCs are Tdtomato+ (25). Similarly, *Rbm24^-/-^* HCs were obtained by using the *Atoh1^P2A-Cre/P2A-Tdtomato^; Rbm24^flox/flox^* mice. *Atoh1^P2A-Cre/+^* is an effective Cre driver and has been used in deleting HC genes in our previous studies (20, 26). Those cells were subject to single-cell isolation and full length smart-seq based single cell RNA-seq (27). Briefly, 30 Ctrl_HCs and 87 *Rbm24^-/-^* HCs were manually picked (Figure 1A). Four cell clusters were formed when the total 117 cells were mixed (Figure 1B), which matched the 4 different cell types: Ctrl_OHCs (16, cell number), Ctrl_IHCs (14, cell number), *Rbm24^-/-^* OHCs (52, cell number) and *Rbm24^-/-^*IHCs (35, cell number), as illustrated in Figure 1C. Note that those HCs were defined as IHCs or OHCs according to IHC specific marker *Tbx2* and *Slc17a8*, and OHC specific gene *Bcl11b* (Figure 1D). We confirmed that *Rbm24* was highly expressed in Ctrl_IHCs and OHCs but was undetectable in all *Rbm24^-/-^* IHCs and OHCs (Figure 1D). The differentially expressed genes (DEGs) between WT and *Rbm24^-/-^* OHCs as well as between WT and *Rbm24^-/-^* IHCs were calculated, respectively (Volcano maps) (Figure 1E). Genes involved in cilium organization were the most enriched in the DEGs that were up-regulated in *Rbm24^-/-^*IHCs and/or OHCs, including *Rfx1, Ift88 and Foxj1* (Supplemental Figure 1). It is known that *Rfx* gene family and *Ifn88* are involved in mouse hair bundle development (28, 29), and *Foxj1* is also important in vertebrate cilia biogenesis (30, 31). The list of all DEGs and Gene Ontology (GO) term genes were included in Supplemental table 1.

**Figure 1.**
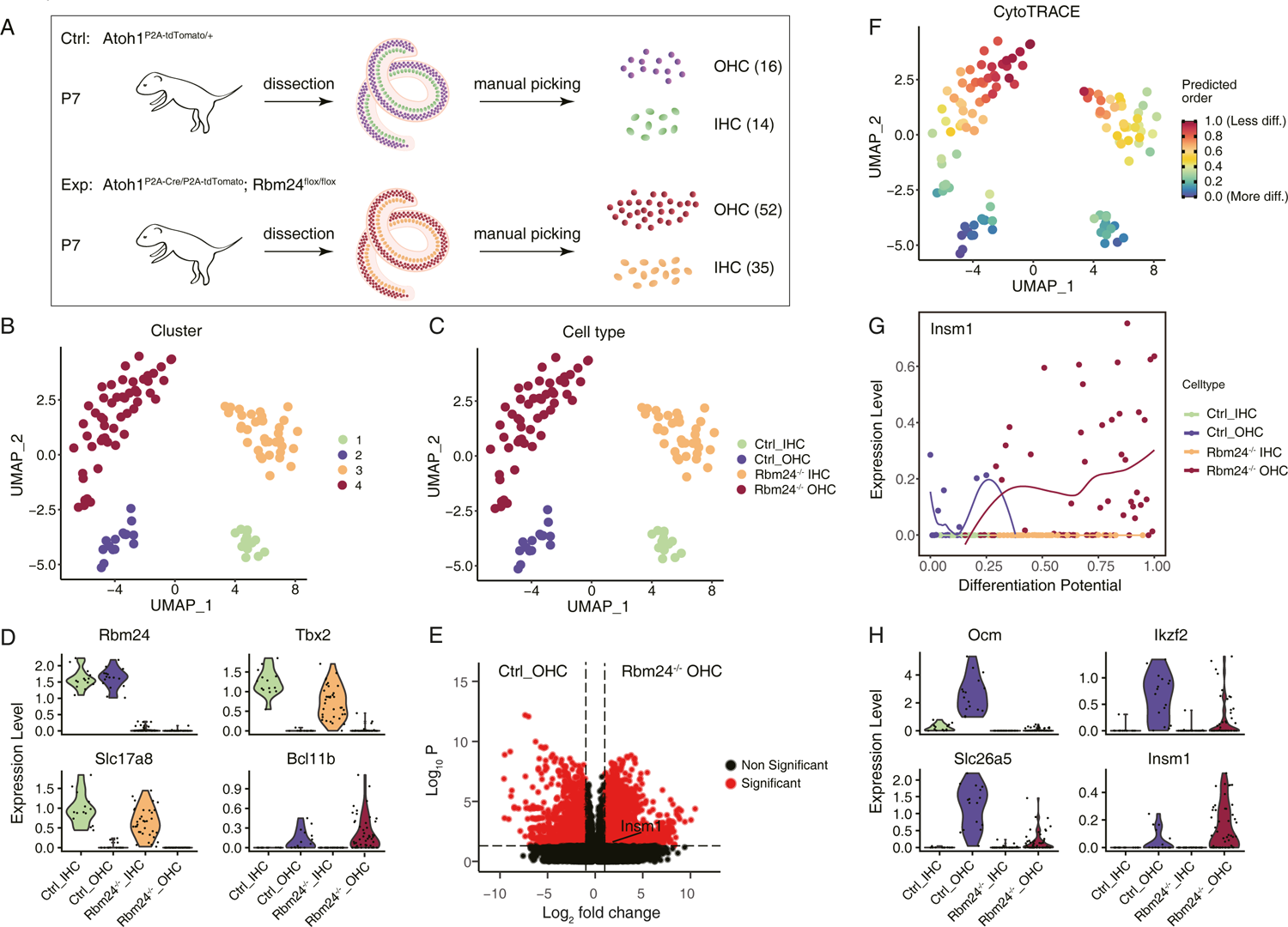
Single-cell transcriptomic analysis of control and *Rbm24^-/-^* HCs. **(A)** An illustration of how we manually pick control IHCs and OHCs, and *Rbm24^-/-^*IHCs and OHCs at P7, respectively. **(B-C)** All picked HCs are subject to unsupervised UMAP analysis and four cell clusters are formed (B), which exactly match each of the 4 cell types in (C). **(D)** Violin plot of four genes, *Rbm24, Tbx2, Slc17a8* and *Bcl11b*. Rbm24 is only detected in WT HCs, and Tbx2 and Slc17a8 are only detected in IHCs, whereas Bcl11b is only detected in OHCs. **(E)** Volcano plot of differentially expressed genes between Ctrl and *Rbm24^-/-^*OHCs. Insm1 is specifically highlighted. **(F-G)** CytoTRACE analysis of the picked HCs. The *Rbm24^-/-^* HCs in general are in the less differentiated status, compared to the Ctrl HCs (F). The expression level of Insm1 in the *Rbm24^-/-^* HCs is higher in the Ctrl HCs (G). **(H)** Violin plot of four OHC genes, *Ocm, Ikzf2, Slc26a5,* and *Insm1.* Opposed to *Insm1, Ocm, Ikzf2, Slc26a5* are expressed in a lower level in *Rbm24^-/-^* OHCs than in Ctrl OHCs.

Moreover, cytotrace analysis, which is widely used to assess the differentiation status (32), was applied to determine whether the global HC differentiation program was altered upon *Rbm24* deletion. It showed that both *Rbm24^-/-^* IHCs and *Rbm24^-/-^* OHCs seemed less differentiated than the WT counterparts (Figure 1F). In this study, we focused on OHCs because *Rbm24^-/-^* OHCs, but not *Rbm24^-/-^* IHCs, are degenerated at P19 (24). The OHC marker Insm1 become undetectable by P2 (19–21). However, the expression level of *Insm1* appeared much higher in *Rbm24^-/-^*OHCs (less differentiated) than in the Ctrl_OHCs (more differentiated) (Figure 1G). Furthermore, violin plot demonstrated that expression levels of other late OHC markers *Ikzf2, Ocm* and *Slc26a5* were lower in *Rbm24^-/-^* OHCs than in Ctrl_OHCs (Figure 1H). It was consistent with the notion that the differentiation status of *Rbm24^-/-^* OHCs was less than WT OHCs, with maintaining early OHC marker Insm1 and not starting expression of *Ikzf2, Ocm* and *Slc26a5* yet at P7.

### Stereocilia development and mechanoelectrical transduction (MET) current are defective in Rbm24^-/-^ OHCs at P3

The morphological change of the stereocilia is a reliable readout to estimate the differentiation status of the hair cells (33). Scanning electron microscope (SEM) assay showed that hair bundle development was defective and the staircase stereocilia was not formed in the *Rbm24^-/-^* OHCs, relative to control OHCs at P3 (Figure 2A and B). We noticed that the morphology of the *Rbm24^-/-^* OHCs resembled the wild type OHCs around P0 (33), again indicating that OHC differentiation was postponed. Our observation was consistent with the notion that Rbm24 is also involved in the hair bundle organization (34, 35). Note that we were puzzled why the cilia development related genes such as *Rfx1, Ift88 and Foxj1* were upregulated in *Rbm24^-/-^* HCs (Supplemental Figure 1). We expected that the cilia organization genes should be down-regulated because their loss-of-function is known to cause defective cilia development (28, 29). Nevertheless, it is possible that both their loss-and-gain of function would cause hair bundle defect, and their maximal expression levels should be negatively controlled by Rbm24.

**Figure 2.**
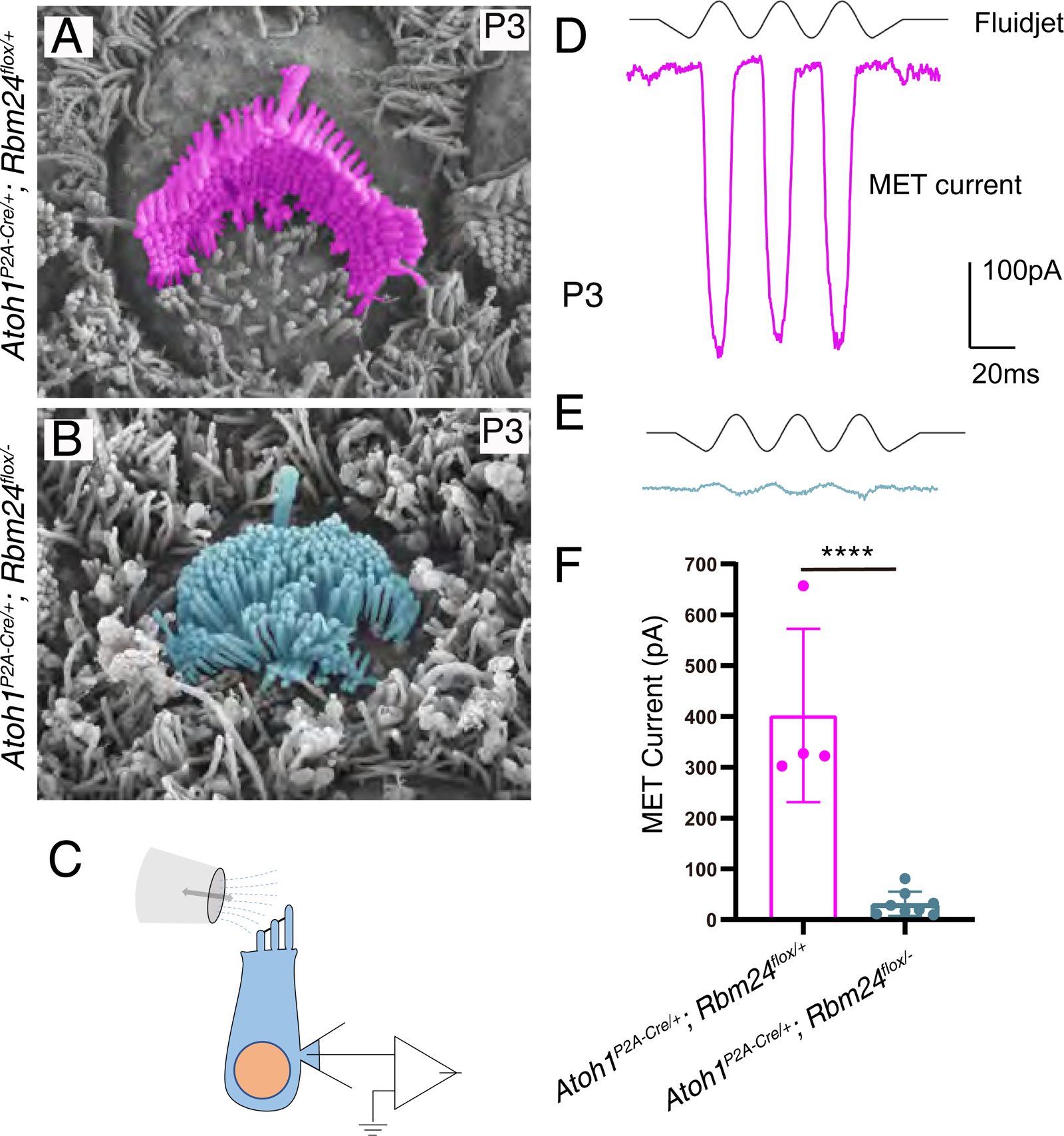
SEM analysis and MET current measurements of Ctrl and *Rbm24^-/-^* OHCs. **(A-B)** Ultrastructure features of hair bundles of OHCs in control *Atoh1^P2A-Cre/+^; Rbm24^flox/+^* (A) and experimental *Atoh1^P2A-Cre/+^; Rbm24^flox/-^* mice (B) at P3. Staircase hair bundles are present in Ctrl OHCs, but not in *Rbm24^-/-^* OHCs. **(C)** A simple illustration of how MET current is measured. **(D-F)** Exampled MET currents in Ctrl (D) and *Rbm24^-/-^* OHCs (E). The amplitude of the MET currents in Ctrl OHCs are much larger than in *Rbm24^-/-^* OHCs. Data are present as Means ± SEM. **** P<0.0001.

To further assess their differentiation status at the electrophysiological level, we further measured the MET currents in Ctrl and *Rbm24^-/-^*OHCs (Figure 2C). We predicted that the amplitude of MET currents would be decreased if loss of Rbm24 delays the OHC differentiation. Large inward current was induced in Ctrl OHCs, however, the amplitude was dramatically diminished in the *Rbm24^-/-^*OHCs at P3 (Figure 3D-F). Collectively, our comprehensive analysis demonstrated that *Rbm24^-/-^* OHCs differentiation was delayed, compared to Ctrl OHCs.

**Figure 3.**
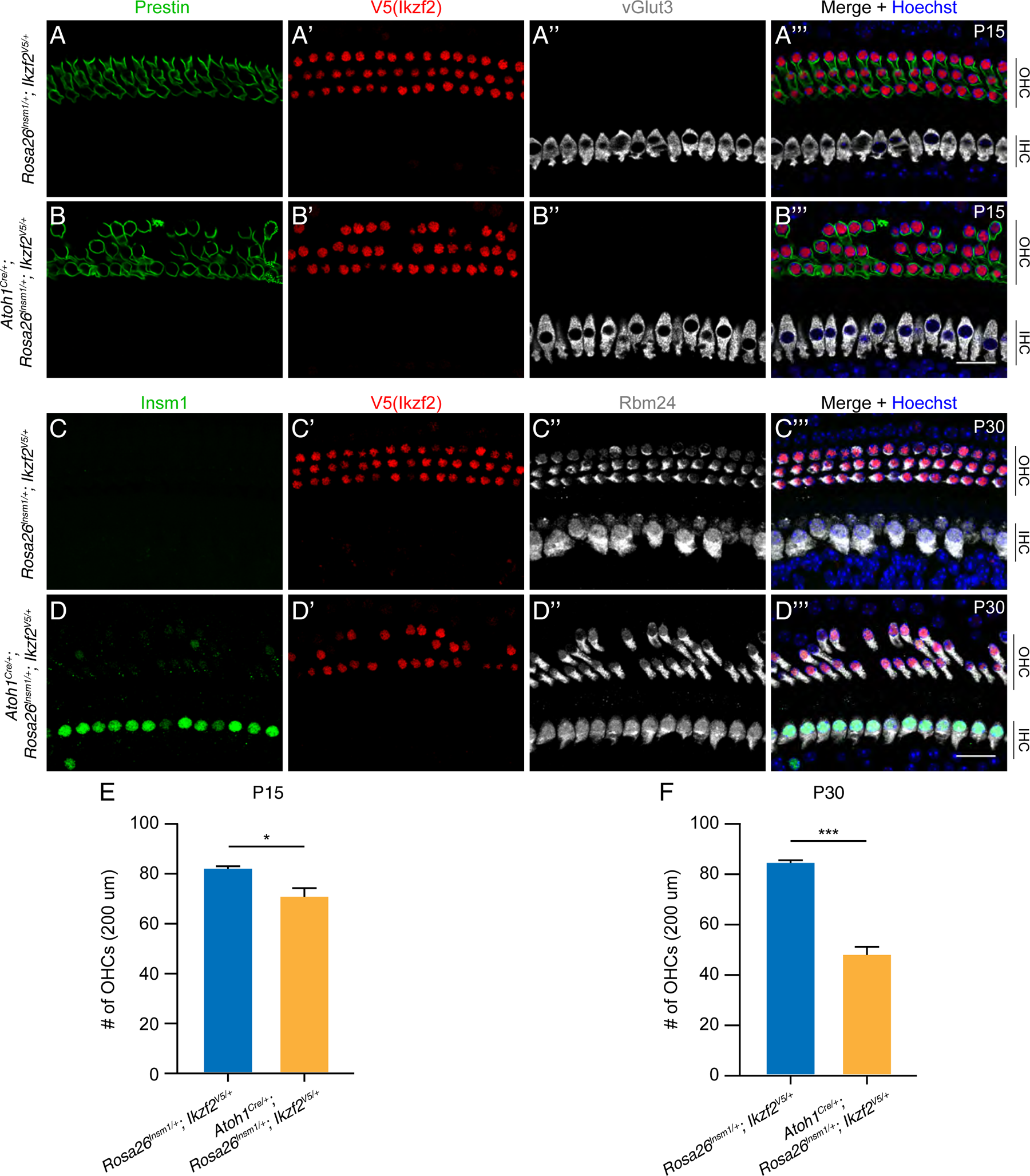
Insm1 overexpression leads to OHC death by P30. (A-B’’’) Triple immunostaining of Prestin, V5 (Ikzf2) and vGlut3 in control *Rosa26^Insm1/+^; Ikzf2^V5/+^* (A-A’’’) and experimental Atoh1^Cre/+^; *Rosa26^Insm1/+^; Ikzf2^V5/+^* mice (B-B’’’) at P15. **(C-D’’’)** Triple labelling of Insm1, V5 (Ikzf2) and Rbm24 in control (C-C’’’) and experimental mice (D-D’’’) at P30. **(E-F)** Quantification of OHC numbers in control and experimental cochleae at P15 (E) and P30 (F). Data are present as Means ± SEM. *p<0.05, *** P<0.001. Significant OHC death occurs in experimental mice at P30. Scale bars: 20 µm (B’’’ and D’’’).

### Insm1 protein expression is prolonged in the *Rbm24^-/-^* OHCs

We next determined whether Insm1 protein expression indeed was reactivated in *Rbm24^-/-^* OHCs at P7. *Atoh1^P2A-Cre/+^; Rbm24^flox/+^* (control group) and *Atoh1^P2A-Cre/+^; Rbm24^flox/-^* (experimental group) mice were characterized in parallel at P7. Rbm24 was expressed in control Pou4f3+ OHCs, but completely disappeared in all Pou4f3+ OHCs of the experimental group mice (Supplemental Figure 2A-B’’’). Insm1 protein was detected in 83.7% ± 5.7%, 67.8% ± 6.4% and 16.1% ± 3.4% of the OHCs in the experimental mice (Supplemental Figure 2C). No OHC death happened in experimental mice by P7. Furthermore, we showed that Insm1 protein was maintained in the Pou4f3+/Rbm24-OHCs at P14 when OHC degeneration had started (arrows in Supplemental Figure 2D-E’’’). On average, there were 21.0 ± 7.6 Insm1+ OHCs per 600 μm in the cochleae of the *Atoh1^P2A-Cre/+^; Rbm24^flox/-^* mice (Supplemental Figure 2F). It suggested that Insm1 protein expression could be maintained in the *Rbm24 ^-/-^*OHCs until they died. Altogether, our Insm1 immunostaining data validated the credit of the single cell RNA-seq of the *Rbm24^-/-^* OHCs and supported that Rbm24 is needed in repressing Insm1 expression in OHCs after P2.

Note that there is a temporal window, between embryonic day 15 (E15) and P2, during which Rbm24 and Insm1 are co-expressed in wild type IHCs and OHCs (21, 23). It remains completely unclear why Insm1 expression is permitted in OHCs between E15 and P2, but not after P2. Further investigations are warranted to fully understand the molecular mechanisms underlying many temporally expressed genes in OHCs, including *Atoh1, Insm1* and *Bcl11b*, which would undoubtedly help us to deeply understand the precisely regulated HC differentiation program.

### Insm1 overexpression also results in OHC death

To determine whether the prolonged expression of Insm1 is one of the main causes that account for cell death of *Rbm24^-/-^* OHCs, we used the *Atoh1^Cre/+^* driver to induce ectopic Insm1 expression in OHCs by using the *Rosa26*-CAG-loxp-stop-loxp-Insm1-P2A-Tdtomato/+ (*Rosa26^Insm1/+^* in short) mice strain (20). Here, we chose *Atoh1^Cre/+^* as the driver because its Cre activity is lower and targets much less SCs than the *Atoh1^P2A-Cre/+^* (26, 36). It is known that OHCs overexpressing Insm1 are intact by P7 (20). However, compared to the OHCs in control *Rosa26^Insm1/+^*; *Ikzf2^V5/+^*mice (Figure 3A-A’’’ and Figure 3C-C’’’), mild OHC death happened at P15 (Figure 3B-B’’’ and Figure 3E), but significant OHC death occurred at P30 in *Atoh1^Cre/+^; Rosa26^Insm1/+^*; *Ikzf2^V5/+^* (Figure 3D-D’’’). There were 83.8 ± 1.1 OHCs per 200 μm cochleae of control mice, in contrast, only 47.3 ± 3.2 OHCs existed in *Atoh1^Cre/+^; Rosa26^Insm1/+^*; *Ikzf2^V5/+^*mice at P30 (Figure 3F).

Collectively, our data demonstrated persistent Insm1 expression was detrimental to OHCs. Thus, the prolonged Insm1 protein expression in OHCs in the absence of Rbm24 should be one of key reasons to explain why *Rbm24^-/-^* OHCs eventually degenerate. Notably, persistent expression of Atoh1, which is also transiently expressed in HCs (26), also cause HC death at adult ages (37). It remains unclear whether persistent Atoh1 and Insm1 expression share the same molecular mechanism underlying HC death.

### The phenotype of OHC degeneration is alleviated in the absence of both Rbm24 and Insm1

If Insm1 reactivation is the key accounting for cell death of *Rbm24^-/-^*OHCs, OHC death should be markedly mitigated when both *Rbm24* and *Insm1* are inactivated simultaneously. Four different mouse models were characterized at P19 (Supplemental Figure 3A-D’’’): 1) *Atoh1^P2A-Cre/+^; Rbm24^flox/+^; Insm1^flox/+^* (control), 2) *Atoh1^P2A-Cre/+^; Insm1^flox/LacZ^*, 3) *Atoh1^P2A-Cre/+^; Rbm24^flox/-^*, and 4) *Atoh1^P2A-Cre/+^; Rbm24^flox/-^; Insm1^flox/LacZ^*. *Insm1^LacZ/+^* is a null allele and LacZ replaces the Insm1 coding sequences (38). Compared to control mice (Figure 4A-A’’’ and Supplemental Figure 3A-A’’’), the phenotype of OHC to IHC fate conversion was observed as expected in the *Atoh1^P2A-Cre/+^; Insm1^flox/LacZ^* mice at P19 (Supplemental Figure 3B-B’’’). However, regardless of the cell fates, the total number (79.0 ± 0.4) of OHCs and IHC-like cells in the OHC regions per 200 μm in *Atoh1^P2A-Cre/+^; Insm1^flox/LacZ^* mice was similar to that (81.5 ± 0.9) in the control mice. Nevertheless, the total cell numbers were 732.7 ± 133.5 in the OHC regions of *Atoh1^P2A-Cre/+^; Rbm24^flox/-^; Insm1^flox/LacZ^* mice, which was significantly more than 52.7 ± 29.8 in *Atoh1^P2A-Cre/+^; Rbm24^flox/-^* mice at P19 (Figure 4B-D). Thus, our data showed that the OHC death was partially alleviated when *Insm1* was further deleted in the *Rbm24^-/-^* OHCs. It further supports that Insm1 reactivation was one of the key contributors to the cell death of *Rbm24^-/-^* OHCs.

**Figure 4.**
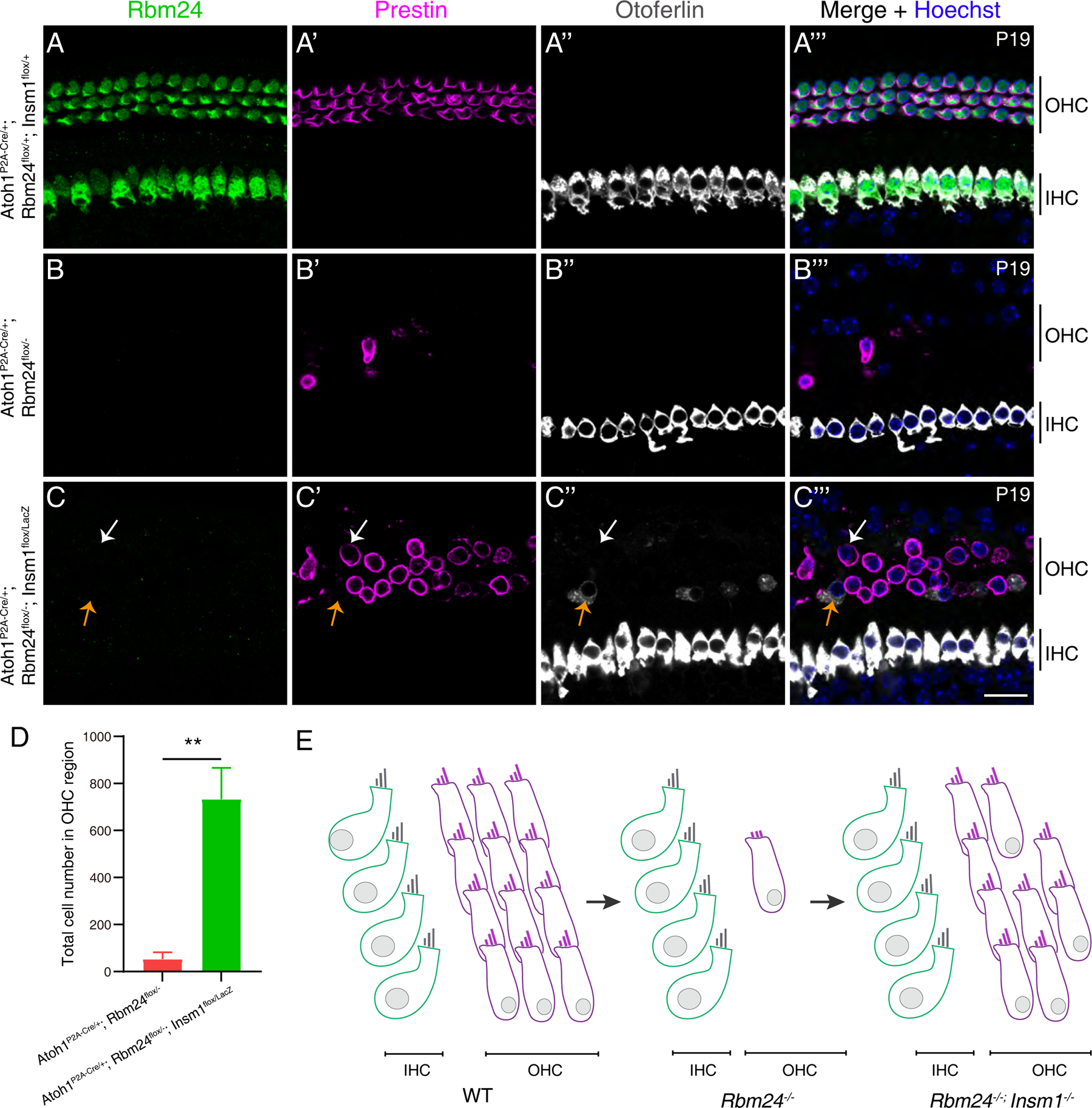
Additional deletion of Insm1 alleviates the cell death of *Rbm24^-/-^* OHCs. (A-C’’’) Triple immunostaining of Rbm24, Prestin and Otoferlin in control *Atoh1^P2A-Cre/+^; Rbm24^flox/+^; Insm1^flox/+^*(A-A’’’), *Atoh1^P2A-Cre/+^; Rbm24^flox/-^* (B-B’’’) and the *Atoh1^P2A-Cre/+^; Rbm24^flox/-^; Insm1^flox/LacZ^* (C-C’’’). White arrows in (C-C’’’) label one Prestin+ OHC without Otoferlin expression, and the orange arrows mark one IHC-like cell that expresses Otoferlin but loses Prestin expression. **(D)** Quantification of the total cell number in the OHC region (either Prestin+ or Otoferlin+) in *Atoh1^P2A-Cre/+^; Rbm24^flox/-^*(red) and the *Atoh1^P2A-Cre/+^; Rbm24^flox/-^; Insm1^flox/LacZ^* (green) cochleae. Data are present as Means ± SEM.** p<0.01. **(E)** A simple cartoon to illustrate how additional loss of Insm1 mitigates the cell death of *Rbm24^-/-^*OHCs. Scale bar: 20 µm (C’’’).

Our previous studies show that Rbm24 is positively regulated by Pou4f3 (39). Pou4f3 is another key TF needed for HC survival (40–44). In addition, *Rbm24* is undetectable in the cochleae of *Atoh1^-/-^* mice (45). Because Pou4f3 is a known target regulated by Atoh1 (25, 46), we suspected that the Rbm24 repression in *Atoh1^-/-^* cochleae is mediated by absence of Pou4f3.

This cascaded signaling from Pou4f3→Rbm24→Insm1 assigns Pou4f3 as the most upstream HC survival regulator, and might explain why *Pou4f3^-/-^*HCs die immediately after HCs are born, but *Rbm24^-/-^* OHCs or OHCs overexpressing Insm1 die at much later ages (24). In sum, our data provided new insights into how Rbm24 controls OHC survival and would pave the way for future OHC protection to prevent the age-related hearing loss.

## MATERIALS AND METHODS

### Mouse models

The *Atoh1^P2A-Cre/+^*, *Rbm24^flox/+^*, *Rbm24^+/-^* and *Rosa26^Insm1/+^* strains were described in detail in our previous studies (20, 24, 26). *Atoh1^Cre/+^*strain was kindly proved by Dr. Lin Gan (Augusta University, USA) (36). *Insm1^flox/+^* and *Insm1^LacZ/+^* strains were kindly provided by Dr. Carmen Birchmeier (Max Delbrueck Center for Molecular Medicine, Germany) (38). Both male and female mice were used in this study. All mice were bred and raised in an SPF-level animal room, and all animal procedures were performed according to the guidelines (NA-032-2022) of the IACUC of the Institute of Neuroscience (ION), Center for Excellence in Brain Science and Intelligence Technology, Chinese Academy of Sciences.

### Histology and immunofluorescence assay

After the mice were anesthetized, 1◊phosphate buffer saline (PBS) was used for heart perfusion, followed by fresh 4% paraformaldehyde (PFA). The post-dissected inner ear tissues were post-fixed with fresh 4% PFA overnight at 4℃ and washed three times using PBS. The inner ears were decalcified with 120mM EDTA (Cat#: ST066, Beyotime) at 4℃. The cochleae were divided into three pieces, apical, middle, and basal turns before immunostaining procedure. The following primary antibodies were used in this study: anti-Rbm24 (rabbit, 1:500, 18178-1-AP, Proteintech), anti-Prestin (goat, 1:1000, sc-22692, Santa Cruz), anti-vGlut3 (rabbit, 1:500, 135203, Synaptic Systems), anti-Insm1 (guinea pig, 1:6000, a kind gift from Dr. Carmen Birchmeier), anti-otoferlin (mouse, 1:500, ab53233, abcam), anti-Pou4f3 (mouse, 1:500, sc-81980, Santa Cruz). After removing the primary antibody by washing thrice (10 mins each) in PBST (1◊PBS containing 0.1%Triton X-100), cochlear samples were further incubated with appropriate secondary antibodies for 3-5 hours at room temperature, and then washed three times in PBST, and counterstained with Hoechst 33342 (1:1000, 62249, Thermo Fisher Scientific). The whole mount prepared cochlear samples were mounted with Prolong Gold antifade medium (P36930, Thermo Fisher Scientific) at room temperature for 1-2 min, and then scanned using a Nikon C2 or Nikon NiE-A1 plus confocal microscope. The confocal images were processed using ImageJ software. The detailed immunostaining protocol was described in our previous study (47).

### Single-cell manual picking and smart-seq RNA-seq library preparation

Cochlear samples were dissected out from control *Atoh1^P2A-Tdtomato/+^*and *Atoh1^P2A-Cre/P2A-Tdtomato^; Rbm24^flox/flox^* mice (experimental group) at P7. The sensory epithelium was carefully dissected out and followed by our single-cell suspension preparation protocol (27). The Tdtomato+ HCs from both group mice were manually picked, immediately followed by reverse-transcriptions and cDNA amplification with the smart-seq HT kit (Cat# 634437, Takara). The post-amplified cDNAs (1 ng per sample) were converted to single-cell sequencing library by using the TruePrep DNA Library Prep Kit V2 for Illumina (Cat# TD503-02, Vazyme) and a TruePrep Index Kit V2 for Illumina (Cat# TD202, Vazyme). The final libraries were subject to paired-end sequencing on the Illumina Novaseq platform. On average ∼4G of raw data per library was yielded.

### Bioinformatic analysis

The raw sequencing data were processed by the zUMIs pipeline (48), and digital gene expression (DGE) matrices were generated. The DGE matrices were analyzed using the R package Seurat (v4) (49). Phase scores were calculated using canonical markers with the “CellCycleScoring” function to mitigate the effects of cell cycle heterogeneity on the “ScaleData” function. Principal component analysis (PCA) was performed using the “RunPCA” function, followed by clustering using the “FindClusters” function. To visualize and investigate the datasets, Uniform Manifold Approximation and Projection (UMAP) was implemented using the “RunUMAP” function. Marker genes for each cluster were identified using the “FindAllMarkers” function, with genes featuring a p-value < 0.05 and the absolute value of log_2_FC (fold change) > 1 as marker genes.

The cell differentiation status was calculated by the R package CytoTRACE (32), which can yield a score the differentiation potential of each cell. To visualize gene expression levels, we extracted the “CytoTRACE” term from the CytoTRACE output and combined it with the Seurat object. The code used for this visualization is available at: https://github.com/mana-W/usual/blob/main/script/cytotraceplot.R. Finally, Metascape platform was used to perform Gene Ontology (GO) analysis (50).

### SEM analysis and MET measurement

Cochlear samples were perfused with 0.9% NaCl (Cat#10019318, Sinopharm Chemical Reagent Co, Ltd.) and fixed overnight with 2.5 % glutaraldehyde (Cat# G5882, Sigma-Aldrich) at 4℃ for overnight. The samples were then washed three times with 1◊PBS and decalcified using 10% EDTA (Cat# ST066, Beyotime) for 1 day. Then, cochlear duct was dissected to three turns, followed by fixation for 1h with 1% osmium tetroxide (Cat#18451, TedPella), and further subject to a second fixation with thiocarbohydrazide (Cat#88535, Sigma-Aldrich) for 30 min and for 1h with 1% osmium tetroxide. Next, the samples were dehydrated using a graded ethanol series (30%, 50%, 75%, 80%, 95%, Cat#10009259, Sinopharm Chemical Reagent Co, Ltd) at 4℃. The cochlear samples were dried in a critical point dryer (Model:EM CPD300, Leica), and further treated with a turbomolecular pumped coater (Model:Q150T ES, Quorum). The post-processed cochlear samples were scanned using a field-emmission SEM instrument (Model:Gemini SEM300, Zeiss).

For the MET measurement, the basilar membrane was precisely dissected from P3 cochleae in the solution containing (in mM): 141.7 NaCl, 5.36 KCl, 0.1 CaCl_2_, 1 MgCl_2_, 0.5 MgSO_4_, 3.4 L-glutamine, 10 glucose, and 10 H-HEPES (pH 7.4). Subsequently, the basilar membrane was placed into a new chamber containing the recording solution consisting of (in mM): 144 NaCl, 0.7 NaH_2_PO_4_, 5.8 KCl, 1.3 CaCl_2_, 0.9 MgCl_2_, 5.6 glucose, and 10 H-HEPES (pH 7.4). Patch pipettes were made of borosilicate glass capillaries (BF150-117-10, Sutter) with the resistances of 3-5 MΩ. The pipette solution was composed of (in mM): 140 KCl, 1 MgCl_2_, 0.1 EGTA, 2 MgATP, 0.3 Na_2_GTP, and 10 H-HEPES (pH 7.2). MET currents were induced by a fluid jet from a pipette with a tip diameter of 5–10 µm. Whole-cell patch clamp on OHCs were conducted with a holding potential of -70 mV (Axon Axopatch 700B, Molecular Devices Corp.). Sinusoidal fluid jet stimuli at 40 Hz were produced using a 27-mm-diameter piezoelectric disc.

### Statistic analysis

All cell numbers are presented as means ± SEM, and statistical analyses were used GraphPad Prism version 8.0.2 (GraphPad Software) and performed Student’s *t-tests* with Bonferroni correction.

## ACKNOWLEDGEMENTS

We thank Dr. Qian Hu from the Optical Imaging Facility of the ION for support with image analysis, and Dr. Lin Gan (Augusta University, USA) for kindly providing the *Atoh1^Cre/+^* strain. We thank Dr. Yanqing Zhong, Ms. Qian Yu, Ms. Jingjing Song, Ms. Yingjie An, and Ms. Ruiqi Wang for helping to pick the WT and *Rbm24^-/-^* HCs.

## SUPPLEMENTAL FIGURE LEGENDS

**Figure S1.**
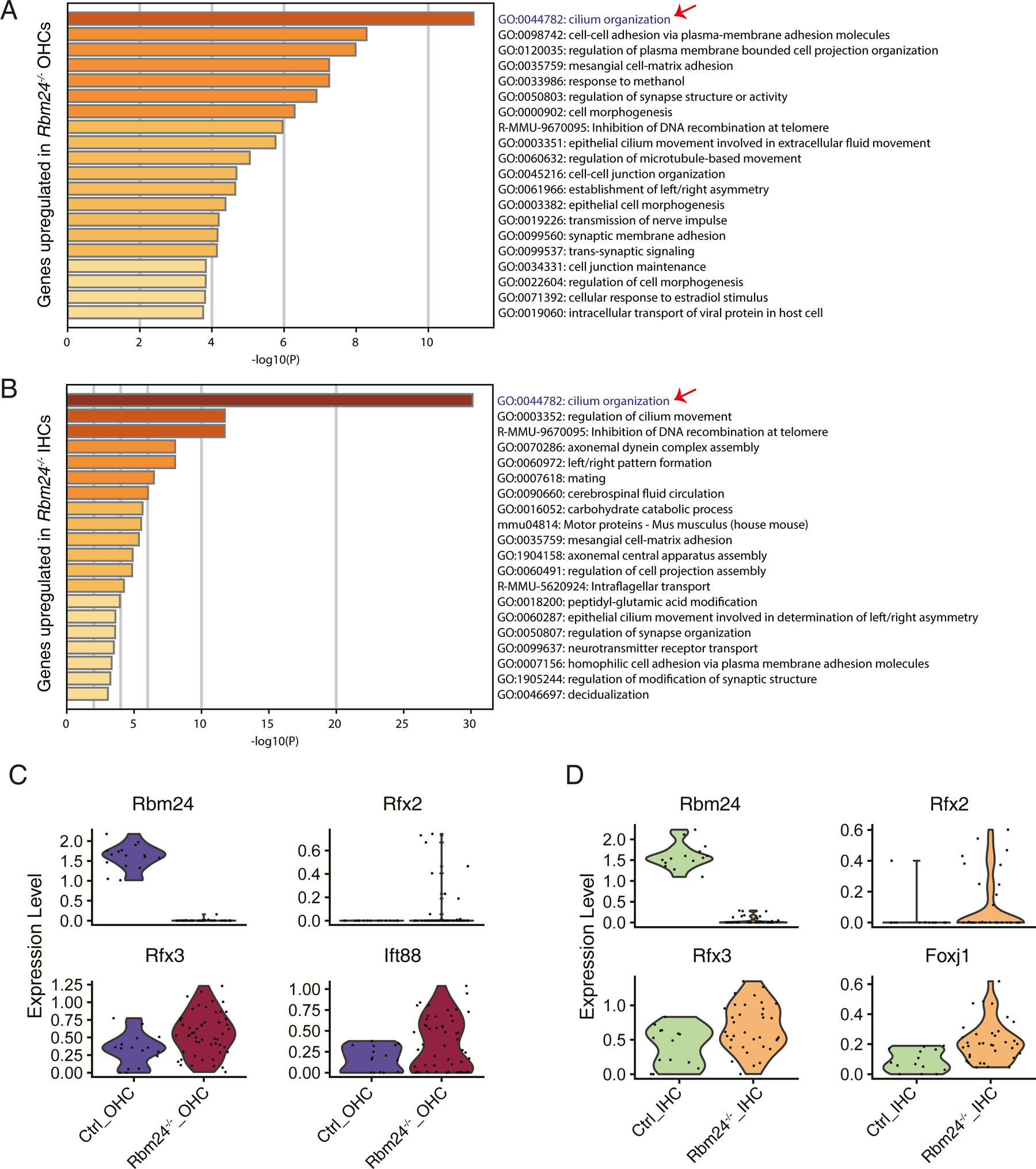
GO analysis of the DEGs between Ctrl and *Rbm24^-/-^* OHCs. **(A-B)** GO terms of the genes that are significantly up-regulated genes in the *Rbm24^-/-^*OHCs (A) or *Rbm24^-/-^* IHCs (B), relative to Ctrl IHCs or OHCs. The red arrows point to the cilium organization related genes that are the most enriched. **(C-D)** Violin plots of *Rbm24, Rfx2, Rfx3*, and *Ift88* between Ctrl and *Rbm24^-/-^* OHCs (C), as well as *Rbm24, Rfx2, Rfx3*, and *Foxj1* between Ctrl and *Rbm24^-/-^*IHCs (D).

**Figure S2.**
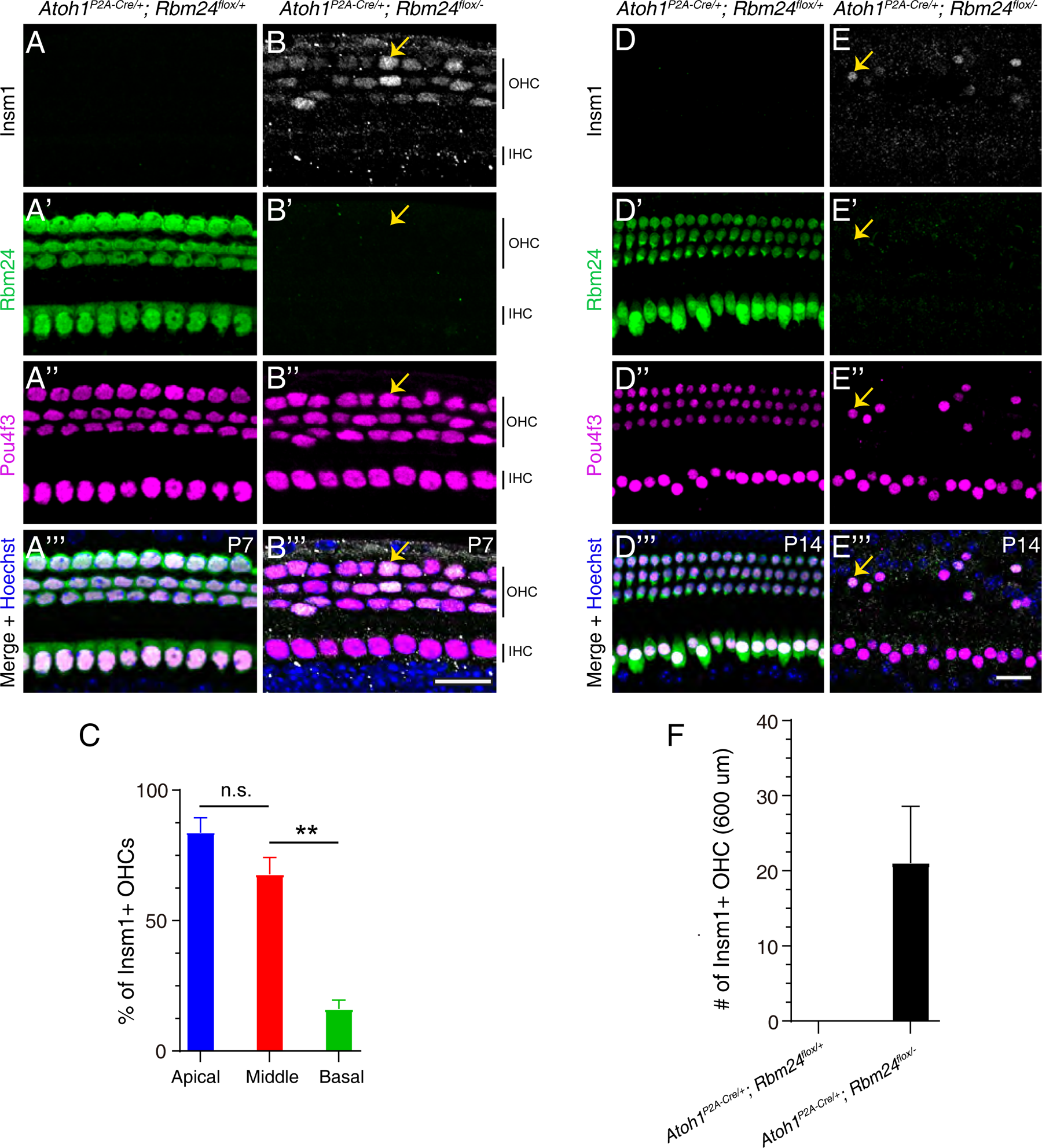
Loss of Rbm24 results in prolonged Insm1 expression in OHCs. (A-B’’’) Triple staining of Insm1, Rbm24, and Pou4f3 in control *Atoh1^P2A-Cre/+^; Rbm24^flox/+^* (A-A’’’) and experimental *Atoh1^P2A-Cre/+^; Rbm24^flox/-^* (B-B’’’) at P7. The orange arrows in (B-B’’’) mark one Insm1+/Pou4f3+ OHC that loses Rbm24. **(C)** Percentages of Insm1+ OHCs in *Atoh1^P2A-Cre/+^; Rbm24^flox/-^* at apical, middle, and basal turns. Data are present as Means ± SEM. n.s.: not significant; ** P<0.01. **(D-E’’’)** Similar immunostaining assay to (A-B’’’) in control (D-D’’’) and experimental (E-E’’’) mice at P14. **(F)** Cell numbers of the Insm1+ OHCs per 600 μm at P14. Again, the orange arrows in (E-E’’’) mark one Insm1+/Pou4f3+ OHC without Rbm24 expression. Data are present as Means ± SEM. Scale bars: 20 µm (B’’’ and E’’’).

**Figure S3.**
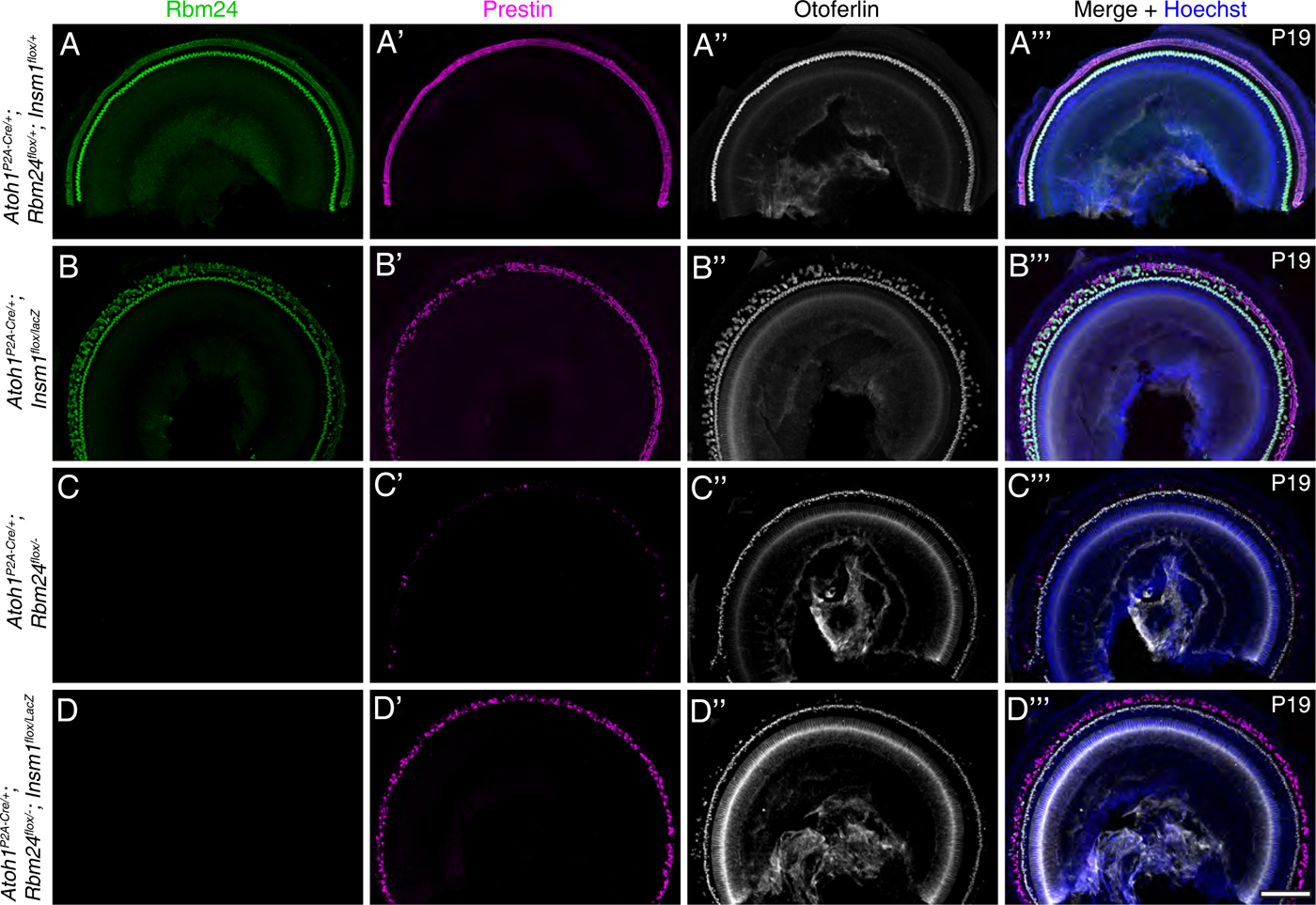
The *Rbm24^-/-^; Insm1^-/-^* OHCs survive longer than the *Rbm24^-/-^*

## REFERENCES

1. Sun Y & Liu Z (2023) Recent advances in molecular studies on cochlear development and regeneration. Curr Opin Neurobiol 81:102745.

2. Luo Z, Zhang J, Qiao L, Lu F, & Liu Z (2021) Mapping Genome-wide Binding Sites of Prox1 in Mouse Cochlea Using the CUT&RUN Approach. Neurosci Bull 37(12):1703–1707.

3. Bermingham NA, et al. (1999) Math1: an essential gene for the generation of inner ear hair cells. Science 284(5421):1837–1841.

4. Zheng J, et al. (2000) Prestin is the motor protein of cochlear outer hair cells. Nature 405(6783):149–155.

5. Liberman MC, et al. (2002) Prestin is required for electromotility of the outer hair cell and for the cochlear amplifier. Nature 419(6904):300–304.

6. Petitpre C, et al. (2018) Neuronal heterogeneity and stereotyped connectivity in the auditory afferent system. Nat Commun 9(1):3691.

7. Shrestha BR, et al. (2018) Sensory Neuron Diversity in the Inner Ear Is Shaped by Activity. Cell 174(5):1229–1246 e1217.

8. Sun S, et al. (2018) Hair Cell Mechanotransduction Regulates Spontaneous Activity and Spiral Ganglion Subtype Specification in the Auditory System. Cell 174(5):1247–1263 e1215.

9. Pan Y, et al. (2023) Fgf8(P2A-3xGFP/+): A New Genetic Mouse Model for Specifically Labeling and Sorting Cochlear Inner Hair Cells. Neurosci Bull 39(12):1762–1774.

10. Roux I, et al. (2006) Otoferlin, defective in a human deafness form, is essential for exocytosis at the auditory ribbon synapse. Cell 127(2):277–289.

11. Jacques BE, Montcouquiol ME, Layman EM, Lewandoski M, & Kelley MW (2007) Fgf8 induces pillar cell fate and regulates cellular patterning in the mammalian cochlea. Development 134(16):3021–3029.

12. Ruel J, et al. (2008) Impairment of SLC17A8 encoding vesicular glutamate transporter-3, VGLUT3, underlies nonsyndromic deafness DFNA25 and inner hair cell dysfunction in null mice. Am J Hum Genet 83(2):278–292.

13. Seal RP, et al. (2008) Sensorineural deafness and seizures in mice lacking vesicular glutamate transporter 3. Neuron 57(2):263–275.

14. Bi Z, et al. (2022) Development and transdifferentiation into inner hair cells require Tbx2. Natl Sci Rev 9(12):nwac156.

15. Garcia-Anoveros J, et al. (2022) Tbx2 is a master regulator of inner versus outer hair cell differentiation. Nature 605(7909):298–303.

16. Kaiser M, et al. (2022) TBX2 specifies and maintains inner hair and supporting cell fate in the Organ of Corti. Nat Commun 13(1):7628.

17. Bi Z, et al. (2024) Revisiting the Potency of Tbx2 Expression in Transforming Outer Hair Cells into Inner Hair Cells at Multiple Ages In Vivo. J Neurosci 44(23).

18. Li X, et al. (2023) In situ regeneration of inner hair cells in the damaged cochlea by temporally regulated co-expression of Atoh1 and Tbx2. Development 150(24).

19. Wiwatpanit T, et al. (2018) Trans-differentiation of outer hair cells into inner hair cells in the absence of INSM1. Nature 563(7733):691–695.

20. Li S, He S, Lu Y, Jia S, & Liu Z (2023) Epistatic genetic interactions between Insm1 and Ikzf2 during cochlear outer hair cell development. Cell Rep 42(5):112504.

21. Lorenzen SM, Duggan A, Osipovich AB, Magnuson MA, & Garcia-Anoveros J (2015) Insm1 promotes neurogenic proliferation in delaminated otic progenitors. Mech Dev 138 Pt 3:233–245.

22. Chessum L, et al. (2018) Helios is a key transcriptional regulator of outer hair cell maturation. Nature 563(7733):696–700.

23. Grifone R, Saquet A, Xu Z, & Shi DL (2018) Expression patterns of Rbm24 in lens, nasal epithelium, and inner ear during mouse embryonic development. Dev Dyn 247(10):1160–1169.

24. Wang G, Li C, He S, & Liu Z (2021) Mosaic CRISPR-stop enables rapid phenotyping of nonsense mutations in essential genes. Development 148(5).

25. Luo Z, et al. (2022) Three distinct Atoh1 enhancers cooperate for sound receptor hair cell development. Proc Natl Acad Sci U S A 119(32):e2119850119.

26. Li S, et al. (2022) Fate-mapping analysis of cochlear cells expressing Atoh1 mRNA via a new Atoh1(3*HA-P2A-Cre) knockin mouse strain. Dev Dyn 251(7):1156–1174.

27. Sun S, et al. (2021) Dual expression of Atoh1 and Ikzf2 promotes transformation of adult cochlear supporting cells into outer hair cells. Elife 10.

28. Elkon R, et al. (2015) RFX transcription factors are essential for hearing in mice. Nat Commun 6:8549.

29. Moon KH, et al. (2020) Dysregulation of sonic hedgehog signaling causes hearing loss in ciliopathy mouse models. Elife 9.

30. Rayamajhi D, et al. (2024) The forkhead transcription factor Foxj1 controls vertebrate olfactory cilia biogenesis and sensory neuron differentiation. PLoS Biol 22(1):e3002468.

31. Stubbs JL, Oishi I, Izpisua Belmonte JC, & Kintner C (2008) The forkhead protein Foxj1 specifies node-like cilia in Xenopus and zebrafish embryos. Nat Genet 40(12):1454–1460.

32. Gulati GS, et al. (2020) Single-cell transcriptional diversity is a hallmark of developmental potential. Science 367(6476):405–411.

33. Sekerkova G, Richter CP, & Bartles JR (2011) Roles of the espin actin-bundling proteins in the morphogenesis and stabilization of hair cell stereocilia revealed in CBA/CaJ congenic jerker mice. PLoS Genet 7(3):e1002032.

34. Wang Y, et al. (2023) RBM24 is required for mouse hair cell development through regulating pre-mRNA alternative splicing and mRNA stability. J Cell Physiol 238(5):1095–1110.

35. Zheng L, et al. (2021) Rbm24 regulates inner-ear-specific alternative splicing and is essential for maintaining auditory and motor coordination. RNA Biol 18(4):468–480.

36. Yang H, Xie X, Deng M, Chen X, & Gan L (2010) Generation and characterization of Atoh1-Cre knock-in mouse line. Genesis 48(6):407–413.

37. Liu Z, et al. (2012) Age-dependent in vivo conversion of mouse cochlear pillar and Deiters’ cells to immature hair cells by Atoh1 ectopic expression. J Neurosci 32(19):6600–6610.

38. Jia SQ, et al. (2015) Insm1 cooperates with Neurod1 and Foxa2 to maintain mature pancreatic β-cell function. Embo J 34(10):1417–1433.

39. Wang G, Gu Y, & Liu Z (2024) Deciphering the genetic interactions between Pou4f3, Gfi1, and Rbm24 in maintaining mouse cochlear hair cell survival. Elife 12.

40. Vahava O, et al. (1998) Mutation in transcription factor POU4F3 associated with inherited progressive hearing loss in humans. Science 279(5358):1950–1954.

41. Masuda M, et al. (2011) Regulation of POU4F3 gene expression in hair cells by 5’ DNA in mice. Neuroscience 197:48–64.

42. Xiang M, et al. (1997) Essential role of POU-domain factor Brn-3c in auditory and vestibular hair cell development. Proc Natl Acad Sci U S A 94(17):9445–9450.

43. Zhu GJ, et al. (2020) Aldh inhibitor restores auditory function in a mouse model of human deafness. PLoS Genet 16(9):e1009040.

44. Erkman L, et al. (1996) Role of transcription factors Brn-3.1 and Brn-3.2 in auditory and visual system development. Nature 381(6583):603–606.

45. Cai T, et al. (2015) Characterization of the transcriptome of nascent hair cells and identification of direct targets of the Atoh1 transcription factor. J Neurosci 35(14):5870–5883.

46. Yu HV, et al. (2021) POU4F3 pioneer activity enables ATOH1 to drive diverse mechanoreceptor differentiation through a feed-forward epigenetic mechanism. Proc Natl Acad Sci U S A 118(29):e2105137118.

47. Liu Z, Owen T, Zhang L, & Zuo J (2010) Dynamic expression pattern of Sonic hedgehog in developing cochlear spiral ganglion neurons. Dev Dyn 239(6):1674–1683.

48. Parekh S, Ziegenhain C, Vieth B, Enard W, & Hellmann I (2018) zUMIs - A fast and flexible pipeline to process RNA sequencing data with UMIs. Gigascience 7(6).

49. Hao YH, et al. (2021) Integrated analysis of multimodal single-cell data. Cell 184(13):3573-+.

50. Zhou YY, et al. (2019) Metascape provides a biologist-oriented resource for the analysis of systems-level datasets. Nature Communications 10.

